# Single cell carbon and nitrogen incorporation and remineralization profiles are uncoupled from phylogenetic groupings of diatom-associated bacteria

**DOI:** 10.1101/2022.07.01.498368

**Authors:** Xavier Mayali, Ty Samo, Jeff Kimbrel, Rhona K. Stuart, Megan Morris, Kristina Rolison, Christina Ramon, Young-Mo Kim, Nathalie Munoz-Munoz, Carrie Nicora, Sam Purvine, Mary Lipton, Peter K. Weber

## Abstract

Bacterial remineralization of algal organic matter is thought to fuel algal growth, but this has not been quantified. Consequently, we cannot currently predict whether some bacterial taxa may provide more remineralized nutrients to algae than others, nor whether this is linked their incorporation. We quantified bacterial incorporation of algal-derived complex dissolved organic C (DOC) and N (DON) and net algal incorporation of remineralized C and N at the single cell level using isotope tracing and NanoSIMS for fifteen bacterial co-cultures growing with the diatom *Phaeodactylum tricornutum*. We found unexpected variability in the net C and N fluxes between algae and bacteria, including non-ubiquitous complex DON utilization and remineralization. We identified three distinct functional categories of metabolic interactions, which we termed macromolecule remineralizers, macromolecule users, and small-molecule users, the latter exhibiting efficient growth under low carbon availability. The functional categories were not linked to phylogeny and could not be elucidated strictly from metabolic capacity as predicted by comparative genomics. Using comparative proteogenomic analyses, we show that a complex DON incorporating strain expressed proteins related to growth and peptide transport, and a non-incorporator prioritized reactive oxygen species scavenging and inorganic nutrient uptake. Our analysis suggests that phylogeny does not predict the extent of algae-bacteria metabolite exchange, and activity-based measurements are indispensable to classify the high diversity of microbes into functional groups. These categorizations are useful for conceptual understanding and mechanistic numerical modeling to ultimately predict the fate of elemental cycles in response to environmental change.

## Introduction

Algal-bacterial interactions are fundamental to primary productivity in the oceans and other surface waters, including algal bioenergy and bioproduct production ponds (Ramanan et al 2016). Studies have shown that bacteria can increase algal productivity by providing vitamins (Croft et al 2005), metal chelators (Amin et al 2009), and algal growth hormones (Amin et al 2015). More fundamentally, bacteria can remineralize nutrients, through, for example, the deamination of organic nitrogen to ammonium (Suleiman et al 2016). It is generally assumed that algae-associated bacteria grow on algal-derived organic matter, making these interactions mutualistic or at least commensal (i.e., the bacteria benefit and the algae are unaffected), but few studies have empirically measured how much C and N bacteria obtain from photosynthetic microalgae. To our knowledge, the reverse process (transfer from bacteria to algae) has never been directly quantified. Measuring exchanges between microalgae and bacteria, that occur between single cells, is critical because when integrated over large volumes, as they have profound consequences for elemental cycling, both in outdoor mass algal cultures and at the global scale (Mayali 2020).

Although direct flux measurements between algae and bacteria are rarely carried out, the last few decades have greatly increased our knowledge of the mechanisms of bacterial C and N recycling in aquatic ecosystems. Algal blooms support a microbial community that progresses over time in response to the changing availability of algal-derived organic matter over the course of weeks (Teeling et al 2012) and as quickly as within one day (Ottesen Elizabeth et al 2014). Generally, algal-associated bacteria that recycle algal organic matter have been divided into those that specialize on polymers and macromolecules, dominated by *Bacteroidia*, and those that specialize on smaller molecules, dominated by *Rhodobacterales* (Buchan et al 2014). In both cases, the primary mechanisms for degradation and uptake involve extracellular enzymes and transporters (Azam et al 1993), and many resources now exist to identify genes involved in such processes, including polysaccharide (Zhang et al 2018) and protein (Rawlings et al 2018) degradation.

Compounds released by bacteria and known to be reincorporated by photoautotrophs for biomass include ammonium (Suleiman et al 2016), amino acids (Liba et al 2006), and carbon dioxide (Danger et al 2007). However, nutrient remineralization has not been identified as a primary mechanism of algal growth promotion by bacteria (Kazamia et al 2016), perhaps because in nature, they are often decoupled in time and space (Cole 1982). Nonetheless, since different bacteria are known to consume different sources of organic matter, it seems plausible that different bacteria likewise express different rates of nutrient remineralization. Thus, there is a need to measure fluxes between algae and bacteria (and bacteria back to algae), identify whether these fluxes are correlated to mutualism, and whether the level of exchange can be predicted by phylogenetic affiliation or genomic content.

A practical approach to quantify metabolic activity and exchange is the use of stable isotope incubations. DNA-stable isotope probing (SIP) was developed to identify organisms that incorporate a pure substrate, for example ^13^C-labeled methanol (Neufeld et al 2008). Further studies have used isotope-labeled extracts from phytoplankton to qualitatively identify bacterial taxa that incorporated these mixed substrates (Nelson and Carlson 2012). A more quantitative analysis from such incubations can be carried out at the single cell level using a nanoSIMS imaging secondary ion mass spectrometer (nanoSIP; Pett-Ridge and Weber 2022). This high-resolution method allows the quantification of isotope incorporation by single cells, including small cells that are attached to one another (Arandia-Gorostidi et al 2017). For example, nanoSIP has been used with mixed bacterial communities growing with one phototrophic alga to show that different bacterial communities have different effects on algal growth and cell-specific carbon fixation, and in some cases this mutualism is mediated by cell-to-cell attachment (Samo et al 2018). However, working with mixed communities makes it challenging to attribute specific impacts to individual taxa, and is also complicated by bacteria-bacteria interactions that might obscure otherwise recognizable interactions. These inherent challenges therefore required us to develop modified culturing and SIP approaches that enable taxon-specific probing and focused observations of bacterial activity and exchange with their algal partner.

Here, we aimed to determine the link between identity and activity of algal-associated bacteria, using isotope tracing to quantify transfer of complex algal C and N to bacteria and remineralized C and N to the algae. We define ‘remineralization’ in these experiments by measurements of net C and N incorporated by the algal cells, so this does not represent gross remineralization of algal organic matter, but only what is made available to the algal cells as they are actively growing, prior to nutrients becoming depleted. We established a culture collection of algal-associated bacteria that was representative of the microbiome associated with an industrially relevant and model alga, *Phaeodactylum tricornutum*. Then, we designed experiments to test two primary hypotheses. Our first hypothesis was that the quantity of complex algal-derived C and N incorporated by algal-associated bacteria correlates with the quantity of remineralized C and N transferred to the algae. Our second hypothesis is that the quantity of algal C and N incorporated and remineralized (i.e. activity) is correlated with bacterial phylogeny and metabolic function as identified by genomic content. Linked activity and genomic potential (Bryson et al 2017) can be useful in order to model complex microbial processes, for example using functional trait approaches (Bouskill et al 2012). To address these hypotheses, we isolated fifteen representative bacteria from mixed community enrichments previously growing with the diatom *P. tricornutum* under autotrophic conditions, with no external sources of organic C. We then reinoculated these bacterial isolates into previously bacteria-free *P. tricornutum* and tested the impact on algal growth, then added isotope-labeled *P. tricornutum*-derived organic matter to these co-cultures to measure organic C and N incorporation by the bacterial cells as well as remineralization of C or N back to *P. tricornutum* cells. We further examined bacterial incorporation of nitrate in the presence of glycolate, a simple C source known to be produced by *P. tricornutum*. In addition to comparative genomics analyses of the 15 strains, we focused on two bacterial isolates representing contrasting phenotypes, linking genotype and phenotype by investigating their most abundantly expressed proteins during growth in algal cultures. These data led us to develop a conceptual framework of bacterial association with microalgae that takes into account nutrient exchange, competition, and metabolism, showing that algal-associated bacteria can be classified into three major functional categories that exhibit distinct strategies allowing them to proliferate in high density algal systems.

## Materials and Methods

### Bacterial genome analysis

Bacterial isolates were cultured from mixed community enrichments grown with previously axenic *P. tricornutum* CCMP 2561 (National Center for Marine Algae and Microbiota; ncma.bigelow.org), as previously reported (Samo et al 2018), and draft genomes were sequenced at the Joint Genome Institute (JGI) through the Community Sequencing Program and annotated and initially analyzed through JGI’s integrated microbial genomes (IMG) pipeline (Chen et al 2019) and dbCAN for carbohydrate active enzyme annotation (Zhang et al 2018). Metabolic pathway annotation for the bacterial genomes was conducted with an approach beyond simple marker gene detection, instead assessing the completeness of pathways where multiple or alternate reactions are involved. We focused on metabolism for the cycling of substrates between algae and bacteria, specifically the pathways for either bacterial utilization of algal-derived compounds or bacterial synthesis of compounds that influence algal physiology (Supplementary Table S1). The database included 359 genes from 55 distinct pathways for biogeochemical reactions related to carbon and nitrogen metabolism, other reactions of primary metabolism (photoheterotrophy, purine and pyrimidine synthesis), secondary metabolism (cell-cell communication and signaling, degradation of aromatic amino acids, vitamin synthesis), and breakdown of polysaccharides and protein. As some genomes were draft quality and not fully complete, we used a threshold of 75 % or greater to consider a pathway as functionally complete.

### Algal growth effects and microscopy

Bacterial influence on algal growth was tested experimentally for each of 15 algal-associated bacterial community isolates as follows. Bacterial isolates were each reinoculated individually with a sterile loop from a single colony into the previously axenic *P. tricornutum* culture and serially transferred monthly for at least 6 transfers before the start of any experiments. Axenic *P. tricornutum* cultures were determined to be bacteria-free via fluorescence microscopy after DAPI staining, as well as sequencing of the 16S rRNA gene (Kimbrel et al 2019), and bacterial abundances and attachment were enumerated from 20-day old, early stationary phase co-cultures, which has been shown to be the growth phase with maximum bacterial attachment (Grossart et al 2005). These co-cultures and the axenic culture were maintained in borosilicate glass 13 mm diameter tubes on a 14/10 light/dark at 75 □mol quanta m^-2^ s^-1^ (cool white fluorescent) at 22 °C in F/2 medium (Guillard 1975) using Instant Ocean salts. We tested growth impacts of co-cultures in 96 well plates (in triplicate) under temperature and nutrient conditions at which they were maintained (22 °C, F/2). Biomass yields were compared to axenic cultures with a one-way ANOVA, adjusting for multiple tests with the Dunn-Sidak method.

### Isotope tracing experiments and nanoSIP analyses

To quantify bacterial remineralization of algal organic matter, we designed a set of experiments where algal-bacterial co-cultures were incubated in the presence of ^13^C and ^15^N labeled algal exudates, obtained through solid phase extraction (Dittmar et al 2008) of axenic *P. tricornutum* spent medium (supplementary information). This labeled material (added at a 2X final concentration to make up for 50 % loss of the extraction protocol) was added to *P. tricornutum* co-cultures (and the axenic culture) in biological triplicates and incubated for 24 hours under a 14/10 light/dark cycle. No-addition and formalin-killed co-culture controls were also included, and for subsequent calculations of substrate incorporation, we quantified the ^13^C and ^15^N enrichment of the added exudate (measured as 18 atom % ^13^C and 45 atom % ^15^N excess) by nanoSIMS after spotting on a Si wafer (Dekas et al 2019). For a second isotope tracing experiment to examine the direct incorporation of nitrate in the presence of a carbon source, we tested the bacterial isolates without *P. tricornutum* cells incubated in unlabeled *P. tricornutum* spent medium. We added 500 □M glycolate (99 % ^13^C, Sigma Isotec) and 800 □M nitrate (98 % ^15^N, Sigma Isotec) for 48 hours to triplicate cultures, which were *P. tricornutum*-bacteria co-cultures filtered through a 0.8 □m syringe filter (so algal cells were removed, but spent medium retained). For a third isotope tracing experiment, we tested the anapleurotic activity for 11 of the 15 strains under extremely low nutrient conditions by incubating them in autotrophic medium (artificial seawater ESAW) with no added carbon plus ^13^C labeled bicarbonate (1.5 mM added) for one week. Samples were prepped and analyzed (see supplemental information) as previously published (Samo et al 2018). Fraction of cellular C and N incorporated from the substrate (C_net_ and N_net_, respectively) was calculated based on the killed controls to represent the initial enrichment of the cells, and the isotope labeling of the added substrates. For the dual-labeled algal organic matter additions, we assumed that the isotope-labeled material was diluted by a 50% addition of unlabeled organic matter (i.e. 66% labeled + 33% unlabeled). We based this calculation on the previous finding that at mid-log growth phase, *P. tricornutum* cultures contain approximately 50 % of the total exuded DOM from a stationary phase culture (López-Sandoval et al 2013). For the nitrate and glycolate experiment, we assumed that unlabeled glycolate and nitrate in the spent medium from early stationary phase was minimal compared to what was added and thus the labeled substrates were calculated to be undiluted. These assumptions were based on previous publications documenting total dissolved organic carbon in spent medium of 300 mM (Chen and Wangersky 1996) and high drawdown of nitrate (Needoba et al 2003) in *P. tricornutum* cultures. Single cell data from biological triplicate incubations were pooled to compare across treatments with a non-parametric Kruskal-Wallis test, followed by a Dunn’s test with multiple comparison p-value adjustment via the Dunn-Sidak method. NanoSIP data from the DOM incubations were used to cluster bacteria into groups using K-means with an optimal cluster count of three as determined by the elbow method in the R factoextra package.

### GC-MS metabolomics and proteomics

The SPE protocol used to extract isotope labeled *P. tricornutum* exudate is known to have low extraction efficiency for small polar metabolites (Johnson et al 2017), and thus our isotope tracing experiments overlooked exchange mediated by these molecules. To examine bacterial incorporation of such small metabolites, we carried out GC-MS analyses of spent media from axenic *P. tricornutum*, axenic *Marinobacter* strain 3-2, media blanks, and *P. tricornutum*-*Marinobacter* co-cultures (all incubated in F/2 media; see supplemental information). We also examined the expression of *Rhodophyticola* 6CLA and *Marinobacter* 3-2 bacterial proteins in the presence of *P. tricornutum* (see supplementary information for method descriptions).

## Results

### Growth impacts, origin, and niche of *P. tricornutum* associated bacteria

Numerically dominant taxa associated with *P. tricornutum*, as measured with cultivation independent 16S rRNA gene sequencing, primarily belong to the Gammaproteobacteria, Alphaproteobacteria, and Bacteroidia (Kimbrel et al 2019). Our isolation efforts yielded strains from 15 unique genera, all belonging to these three taxonomic groups (Table 1), and representing 16S rRNA gene variants identified in outdoor *P. tricornutum* cultures and multi-species laboratory enrichments (Supplementary Fig. S1). Although the culture collection is by no means exhaustive, it spans the major taxonomic groups associated with *P. tricornutum* and provides a practical set of co-cultures that can be tested experimentally to examine patterns of nutrient exchange with their algal host. The majority of the isolates did not impact *P. tricornutum* growth (Table 1, Fig. S2), with the exception of *Marinobacter, Devosia*, and *Alcanivorax*, the presence of which increased algal chlorophyll fluorescence in early stationary growth phase compared to axenic cultures, as previously reported (Chorazyczewski et al 2021). Four bacterial strains were detected attached to *P. tricornutum* cells (*Marinobacter, Oceanicaulis, Roseibium*, and *Yoonia*), all of which originated from phycosphere enrichments where free-living bacteria were washed away (Samo et al 2018). We note that most bacterial cells in these co-cultures were still free-living, with a small percentage detected as algal-attached. We also examined the total cell abundances of the bacterial strains grown in co-culture with no other sources of organic matter besides photosynthetic exudation by *P. tricornutum*. Other than *Henriciella* which was present at such low abundance that it was not able to be counted in our samples (due to *P. tricornutum* cells overloading the filter areas), the bacteria in the co-cultures exhibited abundances ranging from 3×10^4^ to 6×10^6^ cells mL^-1^ (Table 1). There was no statistically significant association between bacterial cell abundances and strain origin (free-living vs attached, or phycosphere enrichments vs algal culture enrichments; data not shown, one-way ANOVA, p>0.05).

**Table 1:** summary of 15 *P. tricornutum* associated bacteria investigated in this study, including enrichment source (free-living or attached), bacterial abundances at stationary algal growth phase, whether co-cultures exhibited some visible attachment to algal cells, and whether algal yield as measured by chlorophyll fluorescence was significantly higher at late log phase compared to axenic cultures; *cultures with two other *Marinobacter* isolates with identical 16S rRNA gene sequence also exhibited significantly higher chlorophyll fluorescence; b.d. = below detection.

| strain | GTDB taxonomy | GB accession | source | abundance | attached | higher yield |
| --- | --- | --- | --- | --- | --- | --- |
| *Henriciella* 6ES | Alphaproteobacteria;Caulobacteriales;Hyphomonadaceae | PRJNA441683 | phycosphere | b.d. | b.d. | no |
| *Oceanicaulis* PT13A | Alphaproteobacteria;Caulobacteriales;Maricaulaceae | PRJNA441690 | phycosphere | 1.58E+06 | yes | no |
| *Tepidicaulis* EA10 | Alphaproteobacteria;Parvibaculales;Parvibaculaceae | PRJNA455451 | free-living | 1.37E+05 | no | no |
| *Devosia* PTEAB7WZ | Alphaproteobacteria;Rhizobiales;Devosiaceae | PRJNA441687 | free-living | 6.84E+04 | no | yes |
| *Roseibium* PT13C1 | Alphaproteobacteria;Rhizobiales;Stappiaceae | PRJNA441686 | phycosphere | 1.33E+06 | yes | no |
| *Stappia* ARW1T | Alphaproteobacteria;Rhizobiales;Stappiaceae | PRJNA441689 | free-living | 2.44E+05 | no | no |
| *Rhodophyticola* 6CLA | Alphaproteobacteria;Rhodobacterales;Rhodobacteraceae | PRJNA441682 | phycosphere | 6.08E+06 | no | no |
| *Yoonia* 4BL | Alphaproteobacteria;Rhodobacterales;Rhodobacteraceae | PRJNA441685 | phycosphere | 1.19E+06 | yes | no |
| *Thalassospira* 13M1/11-3 | Alphaproteobacteria;Rhodospirillales;Thalassospiraceae | PRJNA441688 | phycosphere | 2.05E+05 | no | no |
| *Algoriphagus* ARW1R1 | Bacteroidia;Cytophagales;Cyclobacteriaceae | PRJNA441694 | free-living | 1.96E+06 | no | no |
| *Muricauda* ARW1Y1 | Bacteroidia;Flavobacteriales;Flavobacteriaceae | PRJNA581034 | free-living | 1.95E+06 | no | no |
| *Arenibacter* ARW7G5Y1 | Bacteroidia;Flavobacteriales;Flavobacteriaceae; | PRJNA224116 | free-living | 1.93E+06 | no | no |
| *Pusillimonas* EA3 | Gammaaproteobacteria;Burkholderiales;Burkholderiaceae | PRJNA455452 | free-living | 2.93E+04 | no | no |
| *Alcanivorax* EA2 | Gammaaproteobacteria;Pseudomonadales;Alcanivoracaceae | PRJNA455453 | free-living | 2.05E+05 | no | yes |
| * *Marinobacter* 3-2 | Gammaaproteobacteria;Pseudomonadales;Oleiphilaceae | PRJNA500125 | phycosphere | 1.86E+05 | yes | yes |

### Bacterial incorporation and remineralization of complex algal organic matter

We first aimed to quantify bacterial C and N incorporation of algal exudates during co-cultivation. To address this, we added solid-phase extracted ^13^C and ^15^N labeled extracellular organic matter from axenic *P. tricornutum* to the 15 algae-bacteria co-cultures and used NanoSIMS to quantify net bacterial C and N incorporation after 24 hours. Not including formalin-killed controls, we collected ^13^C and ^15^N incorporation data for a total of 445 bacterial cells (Figs. 1A, S3, S4), finding high variability of incorporation among the isolates.

**Figure 1:**
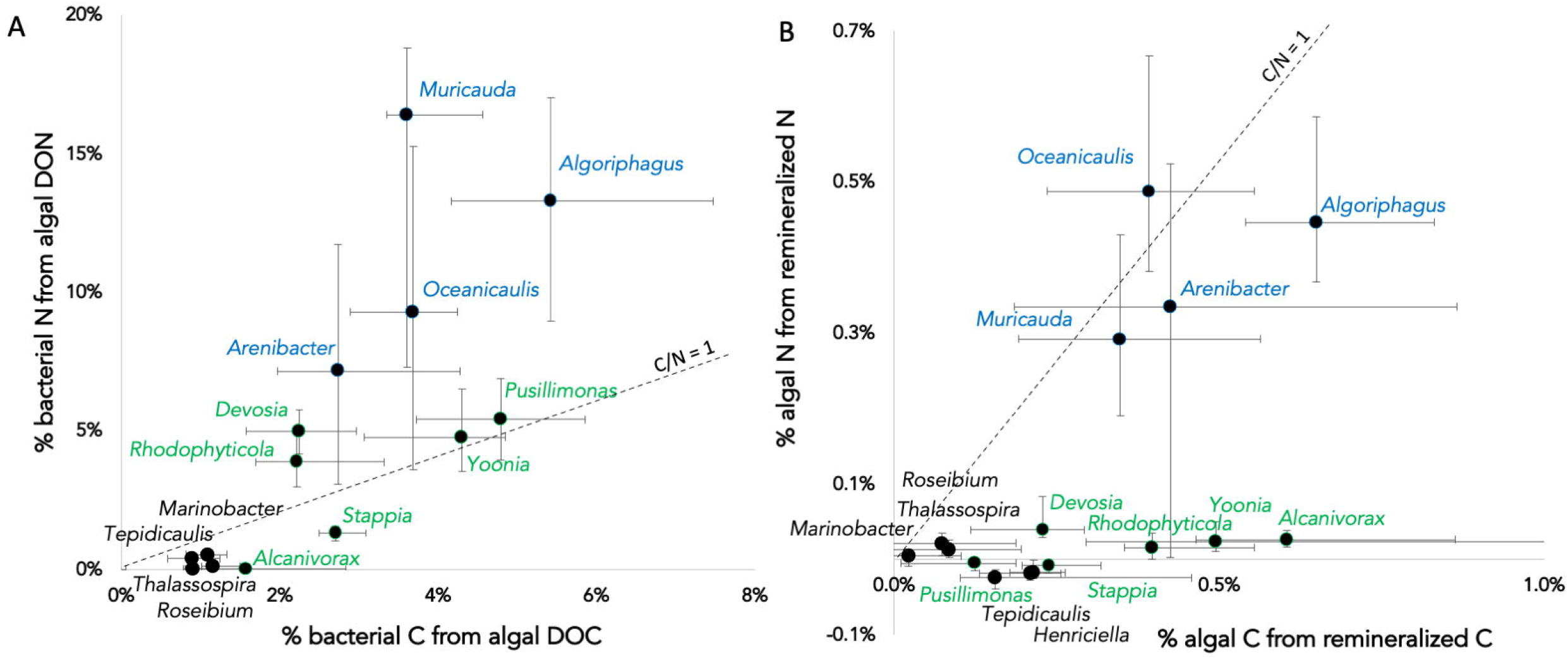
(A) relationship between C and N net incorporation for 15 *Phaeodactylum*-associated bacterial taxa based on single cell NanoSIMS analysis (each point is the population median, error bars represent interquartile range; R^2^ = 0.55). Taxa are colored blue, green, and black based on Kmeans clustering. (B) same relationship for algal incorporation of bacteria-remineralized C and N.

All tested bacteria significantly incorporated algal DON (compared to killed controls) except *Tepidicaulis, Marinobacter*, and *Alcanivorax* (Kruskal-Wallis multiple comparison test, p values 0.2, 0.27, and 0.83, respectively; Fig. 1A). Among the isotopically enriched strains, there was a wide range of incorporation, from median N_net_ values (% of biomass N from the substrate) of 0.4% to 16%. Strains with >5% of their N biomass from added algal DON after 24 hours included *Muricauda, Algoriphagus, Oceanicaulis, Yoonia, Pusillimonas*, and *Arenibacter*. Some of the statistically significantly enriched strains had low N_net_ values (<1%), including *Roseibium* and *Thalassospira*. All bacteria incorporated DOC compared to the killed control (Kruskal-Wallis multiple comparison test; Fig. 1A, S4). Median C_net_ values ranged from 0.9% to 5.4% and bacterial C_net_ and N_net_ were positively correlated (R^2^ = 0.55, p =0.0024, Fig. 1A), suggesting that incorporation of DOC and DON were coupled. The variation in DON/DOC incorporation, however, showed that the bacterial strains exhibited differences in their relative incorporation of algal DON vs DOC. Six strains incorporated low DOC but little or no detectable DON (below the 1:1 line in Fig. 1A): *Stappia, Marinobacter, Roseibium, Thalassospira, Alcanivorax*, and *Tepidicaulis*. On the other end of the spectrum, four strains incorporated more DON relative to DOC: *Muricauda, Arenibacter, Algoriphagus*, and *Oceanicaulis*.

Algal incorporation of labeled C and N also enabled us to quantify remineralized C and N uptake (N = 1149 total algal cells analyzed). Axenic algal cells incorporated low background levels of stable isotopes (not significantly different from the killed controls), so stable isotope enrichment above this background indicated uptake of remineralized DOC and DON. The extent of labeling, as expected for such a relatively short incubation, was low, but detectable for most of the co-cultures. For C, all co-cultures were significantly more enriched than the axenic (Fig. 1B, S4), but many exhibited low levels of enrichment which we do not consider biogeochemically significant. Bacterial DOC incorporation and algal C reincorporated were positively correlated, though just above the 0.05 p-value cutoff for significance (Fig. S5A; regression analysis, R^2^ = 0.27, p = 0.055), illustrating that greater bacterial metabolism of algal DOC generally led to greater C remineralization and subsequent incorporation by the algal cells. Regarding remineralized N, nine of the co-cultures exhibited significantly greater ^15^N enrichment compared to the axenic cultures, but the majority of these were also very low values, except for four: *Algoriphagus* (median N_net_ =0.45%), *Arenibacter* (0.33%), *Muricauda* (0.29%), and *Oceanicaulis* (0.49%; Fig. 1B, S4), which were similar to the levels of DOC incorporation (Fig. 1B). A statistical Kmeans clustering analysis of the DOC and DON incorporation and remineralization data enabled us to assign the 15 bacterial isolates into three functional categories (Fig. S5B), which in two dimensions can be optimally visualized when comparing bacterial ^15^N incorporation and algal ^13^C incorporation (Fig. S5C).

### Bacterial incorporation of low molecular weight C compounds and inorganic N

The SPE protocol, shown to have high extraction efficiency for marine DOM, does not capture all organic compounds, especially small polar molecules such as sugars and amino acids (Johnson et al 2017). Thus, it is likely that components of *P. tricornutum* exudates were not captured by our experiment. To begin to address this issue, we used GC-MS exometabolite analyses to identify small polar metabolites produced by *P. tricornutum* that might be consumed by associated bacteria. For these experiments, we analyzed F/2 media controls, axenic cultures, and co-cultures of *P. tricornutum* and *Marinobacter*. These analyses identified eight small metabolites in greater abundance in *P. tricornutum* cultures compared to the media controls, two of which (glucose and glycolate) were in lower abundance in *P. tricornutum*-*Marinobacter* co-cultures (Supplementary Figure S6; t-test with Dunn-Sidak multiple comparison p-value adjustment). This suggests that *Marinobacter* consumed glucose and glycolate excreted by *P. tricornutum*.

The exometabolite data suggest that *Marinobacter*, and likely other strains that did not incorporate solid phase extracted algal DON, could incorporate small polar C-containing molecules, presumably with nitrate as the N source. We thus tested this hypothesis for all 15 algal-associated strains, using simultaneously added ^13^C labeled glycolate and ^15^N labeled nitrate. For these experiments, we removed *P. tricornutum* cells because they incorporate nitrate and would cross-feed ^15^N to the bacteria via DON release, as shown in the previous experiment. Instead, we included unlabeled *P. tricornutum* spent medium comprised of algal DOC and DON, omitting the SPE. Examining glycolate incorporation, the bacterial strains could be categorized as high incorporators (*Stappia* and *Thalassospira*, with median C_net_ > 5%), medium incorporators (*Marinobacter* with 3% > C_net_ >1.5%), low incorporators (*Alcanivorax, Algoriphagus, Devosia, Yoonia, with* 1% > C_net_ >0%), and non-incorporators (all other strains tested; Fig. S7A). The ^15^N incorporation data further indicate that 9 of the strains incorporated statistically significant levels of nitrate (N_net_ > 0.002). Since isotope labeled nitrate and glycolate were incubated together in excess concentrations, we might expect that their incorporation might be correlated, but nitrate N_net_ was lower than glycolate C_net_ by roughly one order of magnitude (Fig. S7B). This suggests that nitrate incorporation was not enough to fully support glycolate incorporation, and the glycolate-utilizing bacteria incorporated unlabeled DON from the spent media. One of the exceptions was *Algoriphagus*, incorporating the highest levels of nitrate (median N_net_ = 0.87% after 2 days, still relatively low) but very little glycolate (C_net_ = 0.17%). Two other strains (*Muricauda* and *Arenibacter*) did not incorporate glycolate but incorporated significant but again low levels of nitrate. The remaining strains (*Rhodophyticola, Pusillimonas, Roseibium, Tepidicaulis, Oceanicaulis, Henriciella*) did not incorporate nitrate, suggesting they require DON for growth.

The two isotope addition experiments suggested that some of the algal-associated bacteria, somewhat surprisingly, did not incorporate much algal DOM, particularly the third category identified by the Kmeans clustering (colored in black on Fig. 2): *Marinobacter, Henriciella, Roseibium, Thalassospira*, and *Tepidicaulis*. These data led us to develop the hypothesis that these strains are highly efficient at organic matter uptake and may be able to survive on low levels of ambient organic matter. Thus, we attempted to cultivate them on agar plates without any source of organic carbon, and successfully grew *Marinobacter, Roseibium, Thalassospira, Tepidicaulis*, and *Stappia* on artificial seawater agar (Fig. S8; all other strains did not form colonies on such plates). In addition, we tested 11 of the 15 strains in liquid artificial seawater incubated for one week in the presence of ^13^C labeled bicarbonate to quantify activity by measuring anapleurotic carbon fixation, a mechanism that heterotrophs use to replenish lost CO_2_ in the TCA cycle. The bicarbonate uptake experiments in liquid generally matched the results of the growth on artificial seawater agar: efficient strains exhibited detectable DIC incorporation, and non-efficient strains (e.g. *Algoriphagus, Arenibacter, Muricauda, Oceanicaulis*) did not incorporate DIC (Fig. S8).

**Figure 2:**
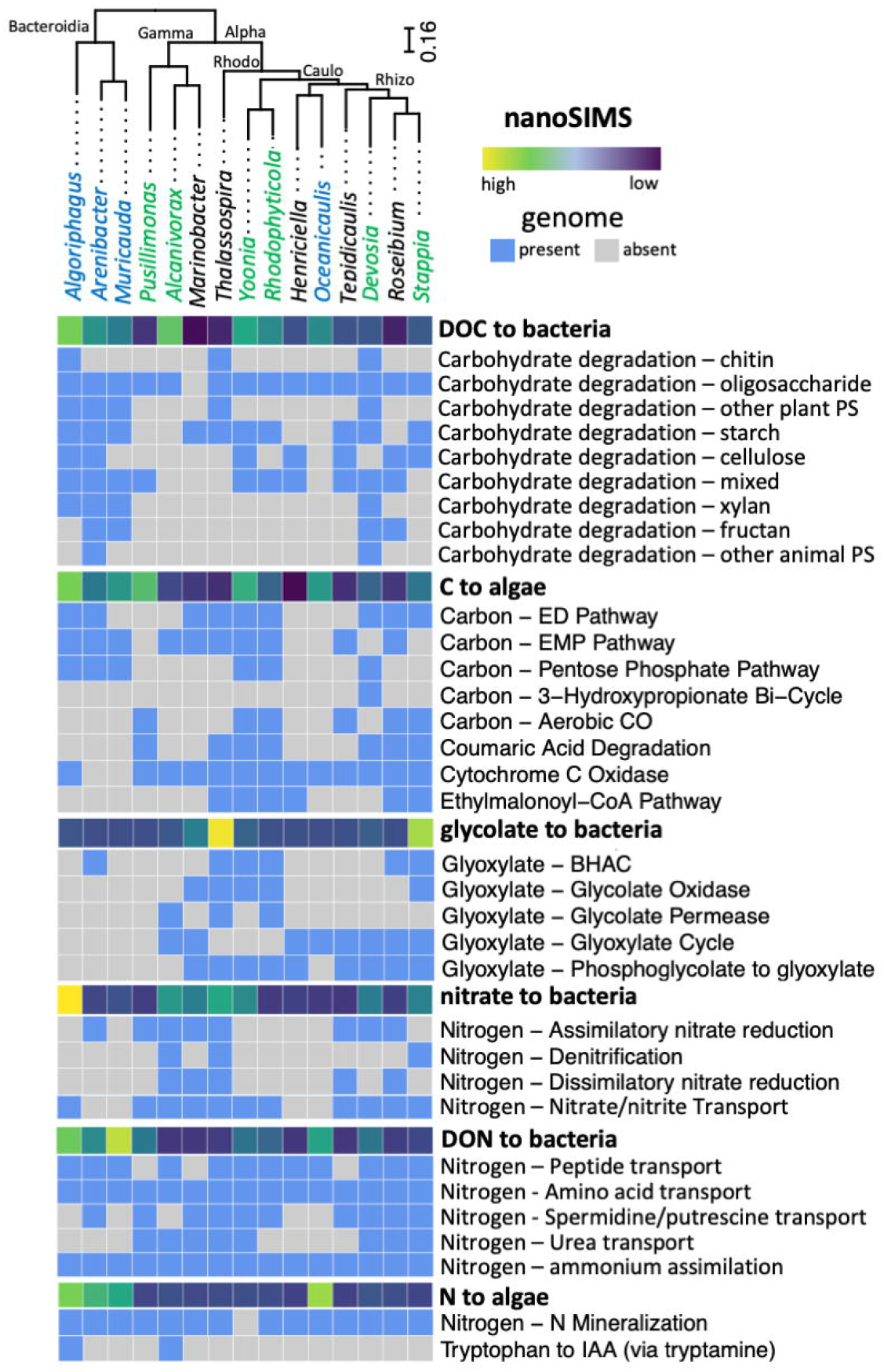
summary of genomic analysis of 15 *P. tricornutum* associated bacteria and comparison with NanoSIMS data (bold). Genus names are colored according to Kmeans clustering from the NanoSIMS data, and bacteria are grouped based on phylogeny of single copy genes

### Genome analyse

Based on cultivation conditions, all 15 bacterial isolates are aerobic heterotrophs, but the genome analysis identified only 12 as being capable of aerobic respiration. The exceptions were the 3 Bacteroidia isolates, suggesting that genome analysis for all metabolic pathways is likely to be biased against this phylogenetic group, and any absence of other pathways within this taxon, and likely other taxa as well, should be interpreted carefully. We briefly summarize these genome analysis results as they relate to the isotope incorporation data, and point the reader to the supplementary information for a longer discussion about other aspects of the genomic capabilities of these organisms.

We first clustered the 15 bacterial strains according to phylogeny (Fig. 2), and mapped isotope incorporation data as well as the presence of metabolic pathways relevant to algae-bacteria interactions. The primary and most evident result of this analysis is that activity as measured by nanoSIMS was not associated with phylogeny. This is most noticeable when the three Kmeans functional categories are mapped onto the phylogeny (Fig. 2), showing little congruence: the functional categories, identified by three different colors, are not organized according to phylogenetic placement. Examining this in more detail, we did not identify links between bacterial incorporation of complex algal carbon and the presence of carbohydrate degradation pathways, nor between carbon remineralization to algae and bacterial central carbon metabolism pathways. We found similar lack of congruence between direct activity measurements and the presence of metabolic pathways with algal organic nitrogen incorporation, nitrogen remineralization, as well as glycolate and nitrate uptake. In most cases (but not always), there was at least one metabolic pathway (or at least a transporter) present in a genome that could explain detected activity by nanoSIMS, but activity could not be predicted by the presence of specific metabolic pathways.

### Protein expression reflects bacterial ecological strategy

We collected proteomics data to examine the expression of bacterial proteins during co-cultivation of *Rhodophyticola* and *Marinobacter* with *P. tricornutum*. We chose to compare these two bacterial co-cultures because they represented distinct bacterial strategies with regards to algal interactions and were grouped into different categories by the Kmeans analysis: *Marinobacter* incorporates glycolate and nitrate but no complex DON, and is an algal-attached mutualist that does not grow to high abundances, whereas *Rhodophyticola* incorporates moderate amounts of complex DON, but not glycolate or nitrate, and is a free-living commensal that grows to high abundances (Supplementary Figure S9). We identified 370 and 160 unique proteins for *Rhodophyticola* and *Marinobacter* co-cultures, respectively, based on our established cutoffs for specificity (Supplementary Table S2). We detected a large number of proteins involved in protein synthesis, transport and energy precursor generation in both taxa (Fig. 3A). We also detected outer membrane proteins in both taxa, and *Marinobacter* expressed outer membrane proteins related to surface attachment (categorized in either outer membrane/cell envelope or motility chemotaxis, Fig. 3A), in agreement with observing *Marinobacter* occasionally attached to *P. tricornutum* cells. *Marinobacter* also highly expressed a greater percentage of stress response proteins, many of which were oxidative stress response proteins, such as superoxide dismutase and catalase-peroxidase KatG, the latter previously associated with bacterial mutualism with a picocyanobacterium (Hennon et al 2018). Under the energy and central metabolism category, both bacteria expressed genes for common pathways such as the TCA cycle, glycolysis, and pyruvate metabolism, but *Rhodophyticola* also highly expressed three syntenous aerobic carbon monoxide (CO) dehydrogenase proteins, suggesting oxidization of CO for energy during co-cultivation. We note that out of the 15 strains, 6 encode this CO oxidation pathway, suggesting this is a common metabolism in co-culture with algae, though measurements of the amount of C processed via CO oxidation are currently lacking. In *Marinobacter*, we detected the narG protein that codes for respiratory nitrate reductase (IMG locus tag Ga0264245_1341), but it was only found in one technical replicate of one of the five biological replicates, so it did not pass our significance threshold. However, it suggests that *Marinobacter* may carry out low levels of denitrification for energy generation.

**Figure 3:**
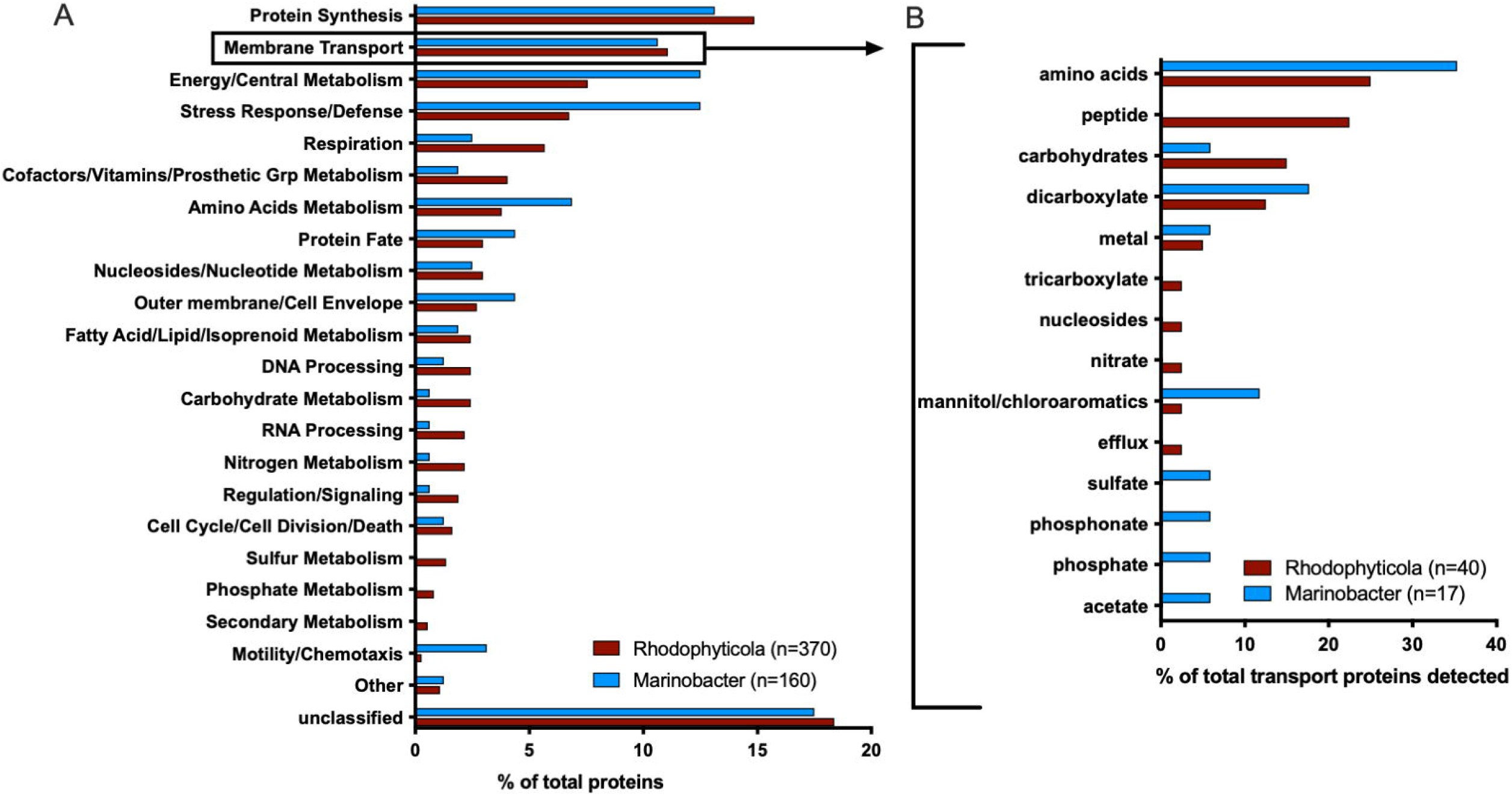
(A) Comparison of functional categories of bacteria-expressed proteins detected from two algal-bacteria co-cultures; (B) further division of annotated transporters expressed at the protein level. Categories based on PATRIC functional classifications to the “Superclass/class” level, including some manual additions to the categories (see supplemental information). “Other” category includes proteins that were less than 1% of the total proteins in both taxa, and include functional classes “clustering based subsystems”, “Iron acquisition and metabolism”, “Metabolite damage and its repair” and “Miscellaneous”.

We examined the transporter protein annotations in more detail as they provide direct links to the nanoSIP incorporation data presented above. These show the two bacteria exhibited differing strategies for organic matter assimilation, with *Marinobacter* expressing proteins for the transport of amino acids, organic and inorganic phosphorus, and acetate, and *Rhodophyticola* expressing proteins for the transport of peptides and small sugars (Fig. 3B). Both strains expressed proteins for transport of dicarboxylate, carbohydrate, and chloroaromatic compounds. These data suggest that *Marinobacter* is adapted to the uptake of small organic compounds, and *Rhodophyticola* for generally larger molecules with more nitrogen content, which agree with the nanoSIP experiments presented above. The lack of ^15^N enrichment in *Marinobacter* cells but their expression of amino acid transporters is also consistent with the extraction protocol being suboptimal for the retention of amino acids.

## Discussion

In this study, we quantified algal DOC and DON uptake and remineralization by bacteria associated with *P. tricornutum*, and used these net flux measurements to categorize bacteria with similar activities into functional categories (Fig. 4). One group, which we call macromolecule remineralizers (identified in blue on Figs. 1, 2, and 4), incorporated high amounts of complex DON (and DOC) and remineralized detectable levels of N to the algal cells. A second group, which we call macromolecule users (green), did not remineralize detectable N to the algal cells. The third group, which we call small-molecule users (black), did not incorporate complex DON, and appeared to be highly efficient recyclers of lost CO_2_ through anapleurotic fixation based on ^13^C bicarbonate incorporation with no algal C present. Within these three groups, there was sometimes wide variation in the levels of C and N incorporation as well as the ability to directly uptake nitrate and glycolate. In addition, there was no pattern regarding attachment or mutualism as they relate to these categories.

**Figure 4:**
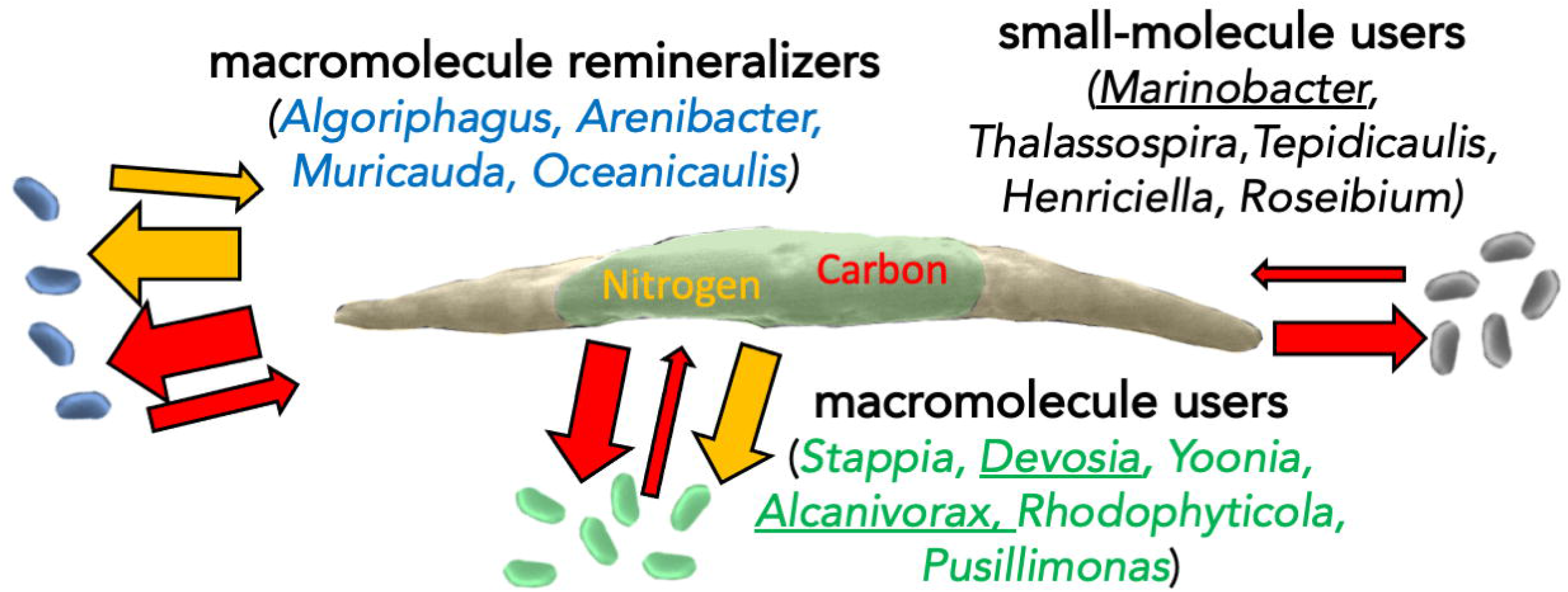
Conceptual figure of bacterial-algal metabolic interactions, based on measurements of C and N incorporation and remineralization by 15 bacterial isolates and remineralization of C and N. Bacteria are classified as macromolecule remineralizers (evidence of nutrient feedback to the algal cells) or users (no N remineralization), and taxa that did not incorporate complex organic N are classified as small-molecule users. Underlined genera have shown evidence of mutualism in co-culture. The thickness of the arrows approximates relative fluxes as measured by isotope incorporation.

Unlike with N, incorporation of DOC and remineralization of C back to algal cells was variable but detectable for all tested co-cultures. The relationship between bacterial C incorporation and remineralized C incorporated by the algal cells in the corresponding co-culture (Fig. S5A) likely reveals the carbon use efficiency of the strains on complex algal-derived organic carbon, with most taxa falling along a positive relationship: greater bacterial C incorporation corresponds to greater remineralized C. Two main exceptions from this trend were *Pusillimonas*, which had very low C remineralization given its C incorporation, and *Alcanivorax*, with the opposite (high C remineralization given its low C incorporation). This suggests that *Pusillimonas* is a highly efficient utilizer of complex algal C, and *Alcanivorax* a highly inefficient one.

Algal incorporation of bacterial remineralized organic matter can sometimes lead to the emergence of mutualism: for example, under long-term cultivation with low nutrient concentrations, picocyanobacterial and heterotrophic bacteria exchange organic carbon for remineralized nutrients (Christie-Oleza et al 2017). Here, we defined mutualism as increased algal biomass yield in co-culture compared to axenic cultures at early stationary growth phase. We did not find that exchange of N was associated with mutualism: none of the three mutualistic strains showed evidence of N mineralization. Furthermore, the highest remineralizers of DON (*Muricauda, Oceanicaulis, Arenibacter*, and *Algoriphagus*) were not mutualistic, and two of those strains (*Muricauda* and *Algoriphagus*) were recently shown to be growth-inhibiting in co-culture with *P. tricornutum* (Chorazyczewski et al 2021). In fact, most of the 15 tested strains, including the three mutualists, provided little to no remineralized N to the algal cells. This suggest that bacterial remineralization of complex algal DON and coupled transfer to the algal cells is not common, at least under the high nitrate concentrations tested here. Bacterial direct nitrate uptake (60% of the isolates) was more prevalent than N remineralization to algal cells, although the data suggest that nitrate did not provide full bacterial N requirement for many of the taxa, based on low nitrate N_net_ compared to glycolate C_net_ (Fig. S7). Bacterial nitrate incorporation for biomass is more energetically expensive than ammonium, and our data confirm that the algal-associated bacteria likely did not use nitrate directly for growth. However, it is possible that algal-associated bacteria used nitrogen cycling for energy. First, some of the strains harbor the full metabolic pathways for dissimilatory nitrate reduction (nitrate to ammonium) and denitrification (nitrate to N_2_O), and strains of *Marinobacter* (Liu et al 2016) and *Thalassospira* (Kodama et al 2008) have been shown to denitrify. Furthermore, some recent evidence suggest that the same *Marinobacter* strain used here positively responds to the availability of F/2 nutrients, likely nitrate, even in the presence of *P. tricornutum* (Kim et al 2021). It remains to be seen whether the mechanism is assimilatory or dissimilatory nitrate reduction, as we did not find proteomic evidence of these processes.

Our data and resulting conceptual framework have implications for better understanding C and N cycling in natural and engineered ecosystems. Due to the complexity of bacterial communities and the difficulty in predicting activity from ‘omics data, the bacterial community is generally considered as a black box with general terms for remineralization, respiration, and other biogeochemical processes (Eichinger et al 2011). Our conceptual framework should help to divide this black box into categories with shared traits, which should then help to more accurately model the impact of different microbial communities on elemental cycling. In particular, the realization that some algal-associated bacteria do not appear to remineralize N, and directly incorporate inorganic N in potential competition with the phytoplankton, should be incorporated into conceptual and eventually numerical models of biogeochemistry.

*Phaeodactylum* is also a model biofuel crop, shown to produce high amounts of biomass and lipids (Wang et al 2020). Understanding the role of the different members of the algal-associated bacterial community could be useful for optimal production of algal biomass or intracellular lipids, reduction of excreted waste DOM, and recycling of nutrients after harvest. One strategy might be to optimize the algal microbiome for different phases of algal cultivation: DOC/DOM remineralizers during early cultivation, and DOC/DOM users post-cultivation, after the algal biomass is harvested, prior to nutrient re-use. Microbiome optimization might also be a critical component of algal cultivation efforts that use recycled nutrients from wastewater or other external sources that includes both organic and inorganic nutrients.

## Supporting information

supplementary methods, results, and figures

## Acknowledgements

Thanks to Arnechia Harper (Georgetown University summer intern) for bacterial microscopy counts. This research was carried out at Lawrence Livermore National Laboratory (LLNL) under Contract DE-AC52-07NA27344, as part of the LLNL Biofuels Science Focus Area FWP SCW1039 supported by the Genome Sciences Program of the U.S. Department of Energy’s Office of Biological and Environmental Research. A portion of this research was performed on a project award (10.46936/sarr.proj.2018.50220/60000027) under the FICUS program and the Joint Genome Institute (JGI) CSP (award 1939) and used resources at the JGI and the Environmental Molecular Sciences Laboratory (EMSL), which are DOE Office of Science User Facilities. Both facilities are sponsored by the Biological and Environmental Research program and operated under Contract Nos. DE-AC02-05CH11231 (JGI) and DE-AC05-76RL01830 (EMSL).

