## supplementary methods, results, and figures for "Single cell carbon and nitrogen incorporation and remineralization profiles are uncoupled from phylogenetic groupings of diatom-associated bacteria"

*Isolation of bacteria*. Samples originated from outdoor algal raceways at Texas A&M Agrilife, which were filtered through a 0.8 micron syringe filter and added to axenic *P. tricornutum* cultures (Samo et al 2018). There algal enrichments were further processed by removing free-living bacteria via sequential centrifugation and washing. Both types of samples were spread onto Marine Agar plates and colonies restreaked and purified. One exception was *Henriciella* 6ES, which cannot grow on agar and was isolated via dilution to extinction.

*Generation of isotope labeled algal organic matter.* We grew axenic *P. tricornutum* in 99 atm % ^15^NO_3_^-^ (Cambridge Isotopes) and ~70 atm % ^13^CO_2_, added as 2 mM sodium bicarbonate (99% ^13^C, Sigma Isotec) into background 1 mM DIC (unlabeled) seawater media, in a sealed glass bottle to avoid further dilution of ^13^C from atmospheric CO_2_. The culture was grown until stationary phase and filtered through a 0.2 μm bottletop filter (Corning) and we used solid phase extraction (SPE) of the labeled filtrate with columns previously shown to extract > 50 % of natural marine organic matter (Dittmar et al 2008). Briefly, the filtered exudate was acidified with hydrochloric acid to pH = 2, run through a Bond Elut PPL column (Agilent), rinsed, and eluted in methanol.

*NanoSIMS analyses.* Samples from isotope labeling experiments were fixed with 2 % formaldehyde and filtered onto 0.2 μm white polycarbonate filters (Whatman Nucleopore, GE Healthcare Life Sciences, Pittsburgh, PA). Filters were washed with 0.2 μm filtered water, air dried and wedges of 1/8 size were cut out of each filter using sterile scissors and adhered to aluminum disks using conductive tabs (#16084-6, Ted Pella, Redding, CA) and sputter coated with ~5 nm of gold. Isotope imaging was performed with a CAMECA NanoSIMS 50 at Lawrence Livermore National Laboratory. The primary ^133^Cs^+^ ion beam was set to 2 pA, corresponding to an approximately 150 nm diameter beam diameter at 16 keV. Rastering was performed over 20 x 20 μm analysis areas with a dwell time of 1 ms pixel^-1^ for 19-30 scans (cycles) and generated images containing 256 x 256 pixels. Sputtering equilibrium at each analysis area was achieved with an initial beam current of 90 pA to a depth of ~60 nm to achieve sputtering equilibrium. After tuning the secondary ion mass spectrometer for mass resolving power of ~7000 (1.5x corrected), secondary electron images and quantitative secondary ion images were simultaneously collected for ^12^C_2_^-^, ^13^C^12^C^-^, ^12^C^14^N^-^, and ^12^C^15^N^-^ on individual electron multipliers in pulse counting mode; ^13^C^12^C^-^/^12^C_2_^-^ = 2 ⋅ ^13^C/^12^C and ^12^C^15^N^-^/^12^C^14^N^-^ = ^15^N/^14^N (Pett-Ridge and Weber 2012). All NanoSIMS datasets were initially processed using L’Image (<http://limagesoftware.net>) to perform dead time and image shift correction of ion image data before creating ^13^C/^12^C and ^15^N/^14^N ratio images, which reflected the level of ^13^C and ^15^N incorporation into biomass. To quantify substrate incorporation, cells were identified based on ^12^C^14^N^-^ images, and regions of interest (ROIs) were drawn manually around each algal cell, making sure to avoid including regions of attached bacterial cells, which in some cases were more highly enriched than the algal cells. ROIs for bacterial cells were drawn separately with the automated algorithm using the ^15^N isotope ratio image. For co-cultures that did not incorporate algal organic matter and thus could not be identified by the ^15^N isotope ratio image, ROIs were drawn manually around free-living (i.e. not associated with algal cells) bacteria-size particles using the ^12^C^14^N^-^ image. The isotopic ratio averages and standard error of the mean for each ROI were calculated from ratios by cycle. The ROI data were exported for further statistical analyses in GraphPad Prism.

*GC-MS metabolomics.* Triplicates from all 4 treatments were grown in 50 % normal salinity (16 g Instant Ocean L^-1^), which increased the detection of metabolites by decreasing salt contamination, collected in early stationary phase (after 10 days of incubation) by filtration through 0.2 μm membrane filters (Corning) and frozen at -20 °C. Spent media samples containing extracellular metabolites were dried down completely in a speed vacuum concentrator (CentriVap, LABCONCO, Kansas City, MO). The dried metabolites were chemically derivatized by two step derivatizations as reported previously (Snijders et al 2016). The derivatized metabolites were analyzed by gas chromatography-mass spectrometry (GC-MS) system (Agilent Technologies, Inc. Santa Clara, CA). All the collected mass spectrometer data files were converted to netCDF format and processed using Metabolite Detector (Hiller et al 2009). All the peaks were matched to PNNL’s augmented version of Agilent Fiehn metabolomics database, which has retention index and fragmented spectra of metabolites and additionally cross-checked with NIST20 and Wiley 11^th^ Edition GC-MS databases. Identification of detected metabolites was validated manually to avoid misidentification of metabolites or false positive and negative errors. Peak area values of identified metabolites were compared among treatments.

*Proteomics.* Three *P. tricornutum* cultures including (1) axenic, (2) co-culture with *Marinobacter* 3-2, and (3) co-culture with *Rhodophyticola* 6CLA were acclimated and maintained for at least 3 rounds of growth in fully defined artificial seawater medium (Berges et al 2001) with modified nutrient levels (1.4 mM NO_3_ and 58.7 µM PO_4_). All cultures were maintained at 22 °C, 75 μmol quanta m^-2^ s^-1^, and a 14/10 light/dark cycle. Cultures were then scaled up to 200 mL volume, grown to late exponential phase and used to inoculate five replicate 500 mL cultures, which were incubated for 7 days (cultures were harvested at day 5). Daily samples were collected for growth and microscopy – fluorescence was measured on 1 mL subsamples, another 1 mL fixed with 4% paraformaldehyde and reserved for bacterial microscopy counts. After 5 days, all cultures were harvested, with all flasks processed in parallel to avoid growth phase differences in expression, reserving 10 mL in the flasks returned to the incubator for a final fluorescence measurement on day 7 to confirm mid-exponential growth phase at time of harvest and track bacterial growth via microscopy on fixed samples. The sampled volume (approximately 485 mL) was centrifuged at 3000 x g for 5 min at 4 °C, pellet resuspended in 4 mL sterile ESAW, aliquoted into two 2.0 mL microcentrifuge tubes, and centrifuged for 5 min at 3000 x g at 4 °C. This second supernatant was decanted and pellets snap frozen in liquid nitrogen and transferred to -80 °C for storage.

Bacterial microscopy counts were conducted by diluting fixed samples 10-fold with 0.2 µm filtered water and filtering on 0.2 µm GTBP filters (MilliporeSigma, Burlington MA), mounting on glass slides with 13 µl slide mountant with 5 µg/µl DAPI (Samo et al 2014) and acquiring 10 images (max intensity z-stack per filter on an inverted fluorescence scope (Dmi6000B, Leica Microsystems, Buffalo Grove IL) with a 100X oil objective, bacteria manually counted on each image, and converted to bacterial cells/mL by multiplying the mean number of cells per field of view by the total filter area divided by the field of view area divided by the volume filtered (Porter and Feig 1980).

Samples were transferred to 2 mL snap-cap centrifuge tubes (Eppendorf, Hamburg, Germany) with 0.1 mm zirconia beads and bead beat in a Bullet Blender (Next Advance, Averill Park, NY) at speed 8 for 3 minutes at 4 °C. After bead beating the lysate was spun into a 15 mL Falcon tube at 2000 xg for 10 minutes at 4 °C. Each sample was transferred to new tubes and a bicinchoninic acid (BCA) assay (Thermo Scientific, Waltham, MA USA) was performed to determine protein concentration. Urea was added to each tube to bring the concentration to 8 M (Sigma-Aldrich, Saint Louis, MO) and dithiothreitol (DTT) was added to the samples at 10 mM. The samples were incubated at 60 °C for 30 minutes with constant shaking at 800 rpm. Samples were then diluted 8-fold for preparation for digestion with 100 mM NH_4_HCO_3_, 1 mM CaCl_2_ and sequencing-grade modified porcine trypsin (Promega, Madison, WI) was added to all protein samples at a 1:50 (w/w) trypsin-to-protein ratio for 3 h at 37 °C. Digested samples were desalted using a 4-probe positive pressure Gilson GX-274 ASPEC™ system (Gilson Inc., Middleton, WI) with Discovery C18 100 mg/1 mL solid phase extraction tubes (Supelco, St. Louis, MO), using the following protocol: 3 mL of methanol was added for conditioning followed by 2 mL of 0.1 % TFA in H_2_O. The samples were then loaded onto each column followed by 4 mL of 95:5: H2O:I, 0.1 % TFA. Samples were eluted with 1 mL 80:20 I:H2O, 0.1 % TFA. The samples were concentrated down to ~100 µL using a Speed Vac and a final BCA was performed to determine the peptide concentration and samples were diluted to 0.1 ug/uL with nanopure water for MS analysis.

A Waters nano-Acquity dual pumping UPLC system (Milford, MA) was configured for on-line trapping of a 5 µL injection at 5 µL/min with reverse-flow elution onto the analytical column at 300 nL/min. Columns were packed in-house using 360 µm o.d. fused silica (Polymicro Technologies Inc., Phoenix, AZ) with 2-mm sol-gel frits for media retention and contained Jupiter C18 media (Phenomenex, Torrence, CA) in 5 µm particle size for the trapping column (150 µm i.d. x 4 cm long) and 3 µm particle size for the analytical column (75 µm i.d. x 70 cm long). Mobile phases consisted of (A) 0.1 % formic acid in water and (B) 0.1 % formic acid in acetonitrile with the following gradient profile (min, %B): 0, 1; 2, 8; 20, 12; 75, 30; 97, 45; 100, 95; 110, 95; 115, 1; 150, 1.

MS analysis was performed using a Q-Exactive HF mass spectrometer (Thermo Scientific, San Jose, CA) outfitted with a home-made nano-electrospray ionization interface. Electrospray emitters were prepared using 150 um o.d. x 20 um i.d. chemically etched fused silica (Kelly et al 2006). The ion transfer tube temperature and spray voltage were 320 ºC and 2.2 kV, respectively. Data were collected for 120 min following a 20 min delay from sample injection. FT-MS spectra were acquired from 350-2000 m/z at a resolution of 30 k (AGC target 1e6) and while the top 12 FT-HCD-MS/MS spectra were acquired in data dependent mode with an isolation window of 2.0 m/z and at a resolution of 15 k (AGC target 1e5) using a normalized collision energy of 30 and a 45 sec exclusion time. For each of the co-cultures there were five biological replicates and two technical replicates for a total of 10 runs per co-culture. Bacterial proteins were not abundant relative to the algal proteins, and there were a few peptides that were detected in either the axenic control culture or the incorrect co-culture (e.g. a *Marinobacter*-predicted peptide in the *Rhodophyticola* co-culture). To account for the low abundance and the non-specificity, we limited our analysis to proteins detected in both technical replicates of at least one of the five biological replicates, and proteins that at least 80% of protein counts were detected in the correct samples, which typically allowed for one peptide detected in one of the 20 non-specific sample runs.

Supplementary genomics result.

Here we present further results of the genomic analysis of the 15 bacterial strains associated with *P. tricornutum* laboratory batch cultures (supplementary table 1). We found no evidence of de-novo net carbon fixation, except that *Devosia* showed capacity for carbon fixation via the 3-hydroxypropionate bi-cycle, which remains to be tested experimentally. None of the strains appear to be capable of methanogenesis where methane is produced via anaerobic respiration, as all lack the two possible pathways. However, 5 strains may be able to produce methane aerobically as a byproduct during conversion of methylphosphonate in the C-P lyase pathway. None of the strains appear to be methanotrophs, as all are lacking the pathway to convert methane to formaldehyde. Four strains appear to be capable of light harvesting mechanisms, either through proteorhodopsin-based phototrophy (*Algoriphagus* and *Muricauda*) or the light-harvesting system II of aerobic anoxygenic phototrophy (AAP; *Roseibium* and *Yoonia*).

We next examined both synthesis and degradation of carbohydrates, which are major components of algal organic matter, both extracellular and serving as intracellular storage compounds, the latter primarily as chrysolaminarin in the case of *P. tricornutum*. All bacterial strains demonstrate the capabilities to metabolize carbohydrates via at minimum one of five pathways: Embden-Meyerhof-Parnas (EMP), phosphorylative or semi-phosphorylative Entner-Doudoroff, or pentose phosphate pathway (oxidative or non-oxidative). Of the five carbohydrate metabolism pathways, only the non-oxidative pentose phosphate pathway was identified across all genomes. An average of at least 3 carbohydrate metabolism pathways (x̄ = 3.26) were complete. The maximum number of complete carbohydrate degradation pathways that a genome contained was 5, with *Algoriphagus*, *Arenibacter*, *Yoonia*, and *Rhodophyticola* demonstrating the most diverse metabolic capacities for carbohydrate metabolism. *Pusillimonas*, *Oceanicaulis*, and *Henriciella* contained the fewest complete pathways for carbohydrate metabolism, with just one complete pathway. Regarding carbohydrate synthesis, the glyoxylate cycle, a modified TCA cycle for biosynthesis of carbohydrates, contains 9 pathways for formation of metabolic intermediates. Twelve of the 15 bacterial genomes contain at least one complete of the 9 pathways, with *Rhodophyticola* encoding the greatest number of pathways (n = 7/9 pathways), and *Muricauda* and *Pusillimonas* lacking any complete pathways within the glyoxylate cycle. Eight of 9 pathways in the cycle were identified in at least one bacterial strain, with just the glyoxylate to pyruvate pathway lacking or incomplete from all genomes. The most conserved pathways across the 15 strains are the phosphoglycolate to glyoxylate (n = 10 genomes) and the ethylmalonoyl-CoA (lower – succinyl-CoA) (n = 8 genomes). Five strains are capable of oxidizing glycolate, a product of algal photorespiration (*Yoonia*, *Marinobacter*, *Stappia*, *Thalassospira*, and *Rhodophyticola*), all of which except incorporated it empirically, but surprisingly fewer strains have annotated genes for transporting glycolate across the cell membrane using glycolate permease (n = 3 strains: *Alcanivorax*, *Thalassospira*, *Rhodophyticola*). This suggest that glycolate transporters are not well identified in the databases.

We also examined the potential for synthesis and catalysis of components outside of central carbon metabolism that have been previously implicated in mutualistic or antagonistic interactions with microalgae, but that we did not test empirically in this study. In addition to being produced by plants and algae (including *P. tricornutum*), phenylacetic acid (PAA) and indole-3-acetic acid (IAA) are plant growth-promoting substrates produced by some groups of bacteria. Ten of the strains are capable of phenylacetic acid (PAA) catabolism (n = 10), with two (*Alcanivorax* and *Algoriphagus*) also capable of synthesizing indole-3-acetic acid (IAA) from tryptophan, which they can accomplish using just one of the five potential pathways via amino acid decarboxylase. Another algal compound of interest, is often produced during algal senescence and inhibits bacteria (Lou et al 2012). Seven of the strains can degrade *p*-coumaric acid (known as pCA) (*Devosia*, *Roseibium*, *Stappia*, *Yoonia*, *Rhodophyticola*, *Thalassospira*, and *Pusillimonas*). This compound is known to stimulate production of algicidal roseobacticides, a class of tropodithietic acid (TDA) produced by the Roseobacter clade, but none of the bacterial genomes, including the 4 isolates belonging to Rhodobacteraceae, contain the metabolic pathways for TDA/roseobacticide synthesis. Two bacterial isolates were found to additionally tolerate antimicrobial pCA via the intermembrane phospholipid transport system for maintaining membrane integrity (Calero et al 2018), including *Marinobacter* and pCa-degrading *Thalassospira*. The two Gammaproteobacteria (*Alcanivorax* and *Marinobacter*), as well as an Alphaproteobacterium (*Thalassospira*), contain the genomic capability for the Gammaproteobacteria-specific Gac/Rsm density-dependent secondary metabolism and carbon storage regulation system (Ferreiro and Gallegos 2021), a regulatory pathway involved in signal transduction, suggesting that these taxa can communicate with bacterial cohorts (e.g. quorum sensing, biofilm formation) or perform biochemical signaling with algae.

We also carried out an extensive analysis of the vitamin production capability of the strains, since vitamin exchange is one of the identified mechanisms of algal-bacterial mutualism. Note that the F/2 algal medium contains vitamins B1, B7, and B12. Eleven of the 15 isolates are capable of synthesizing at least one class of B vitamins, with five (*Alcanivorax, Arenibacter, Marinobacter, Pusillimonas, Tepidicaulis*) capable of synthesizing vitamin B7 (biotin) and seven containing at least one complete of three pathways for B12 synthesis (*Alcanivorax*, *Devosia*, *Roseibium*, *Yoonia*, *Stappia*, *Thalassospira*, and *Rhodophyticola*). None of the bacterial strains can synthesize vitamin B1 (thiamine). Vitamin synthesis Of B12 and B7 had no overlap, meaning that if a strain could synthesize B12, it could not synthesize B7, and vice-versa.

Supplementary Figures

Figure S1: Phylogenetic tree of 1,443 16S rRNA gene ASVs from P. tricornutum cultures from Kimbrel et al. 2019, with the branch tips for the 15 isolates highlighted in red.

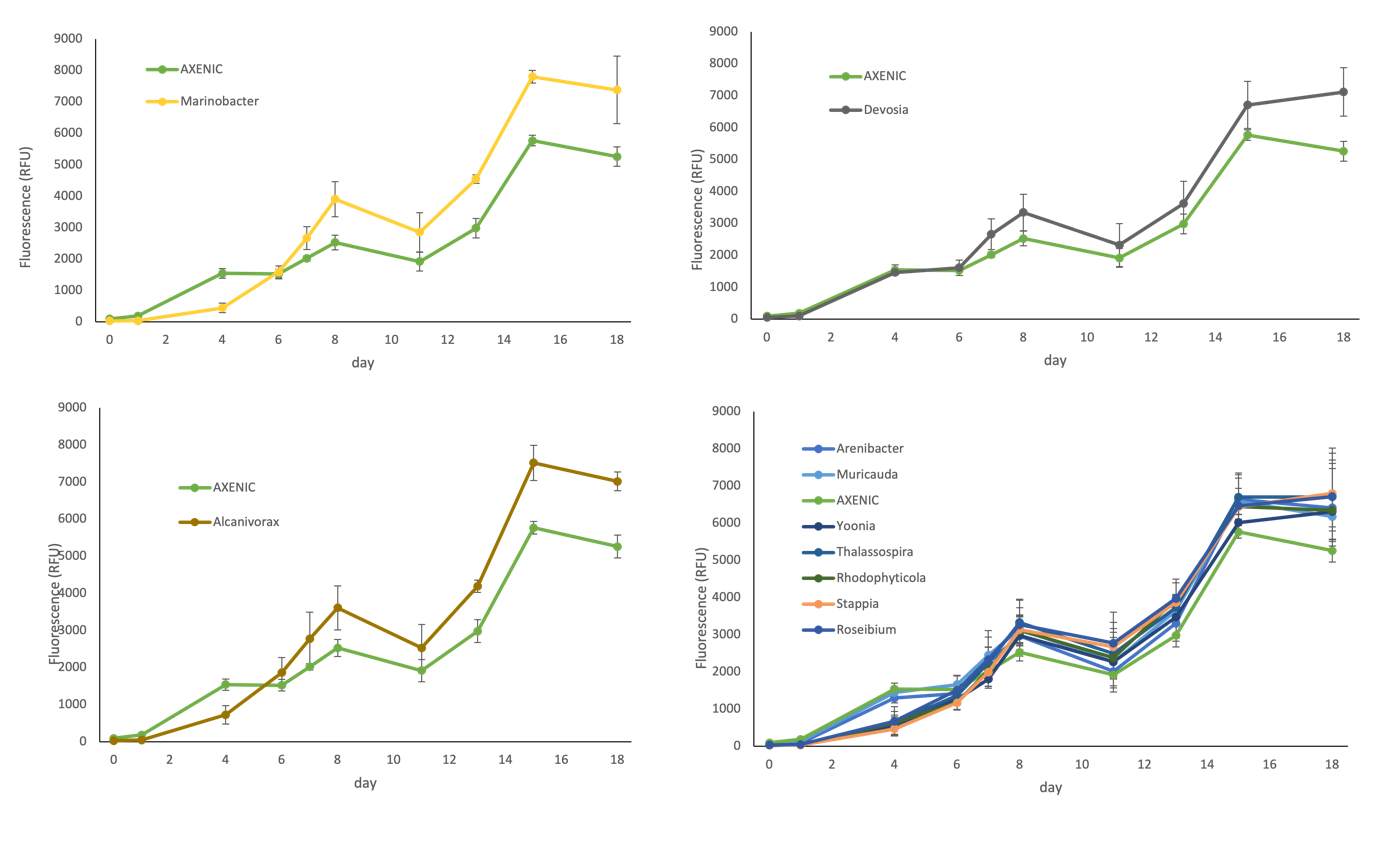

Figure S2 Growth of 3 co-cultures showing increased yield compared to axenic cultures (triplicate, 200 microliter volume, mean +/- standard deviation), and 7 other co-cultures showing equivalent growth to the axenic

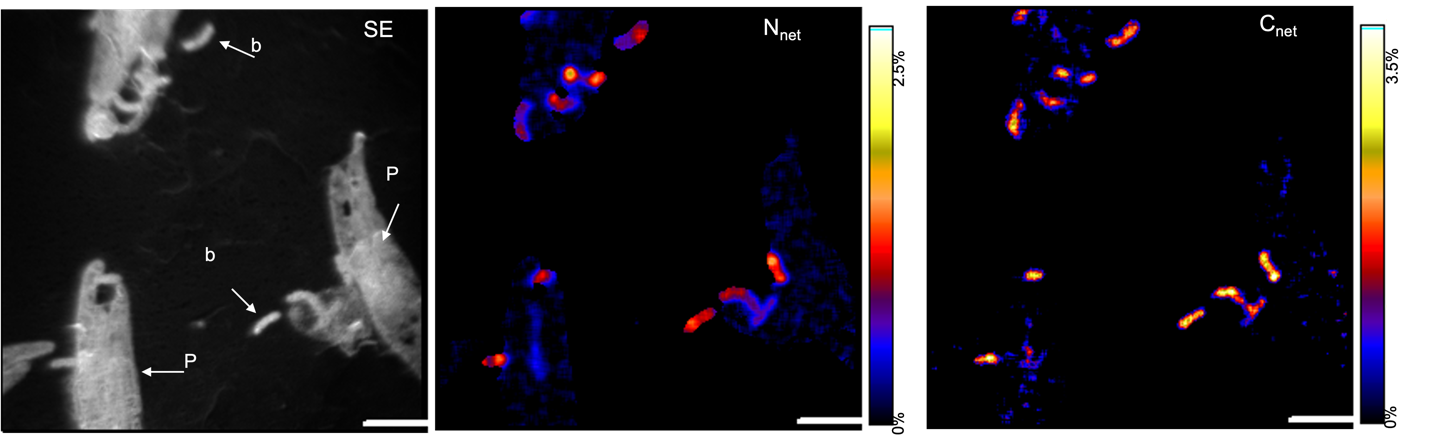

Figure S3: representative NanoSIMS images (secondary electron, SE, and ^15^N and ^13^C enrichment) showing P. tricornutum (P) and bacteria (b) scale bar = 3 microns;

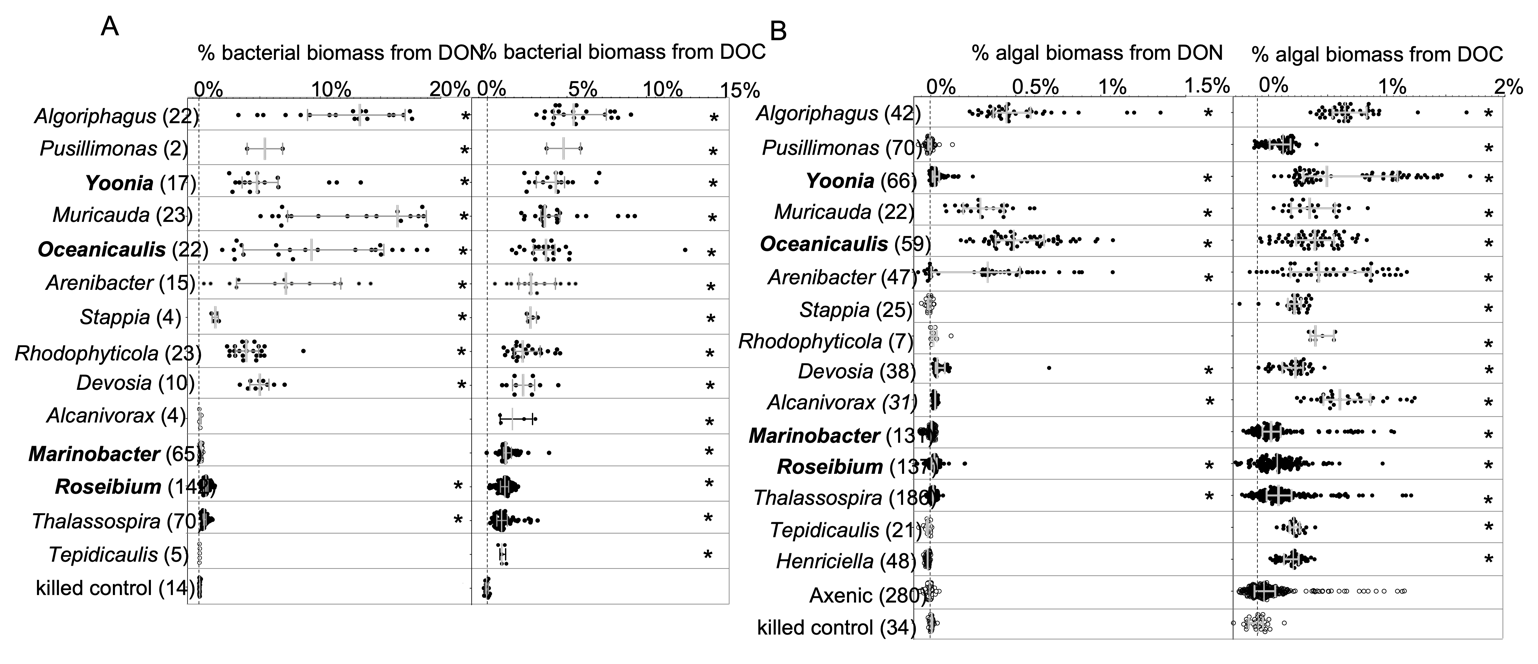

Figure S4: percentage of C and N biomass from algal DOC and DON incorporated (A) by single bacteria (median +/- 95% C.I.) and (B) by single algae as measured by 13C and 15N incorporation (asterisks show cultured significantly more enriched than killed controls; bold name indicates this bacterium can attach to diatom cells);

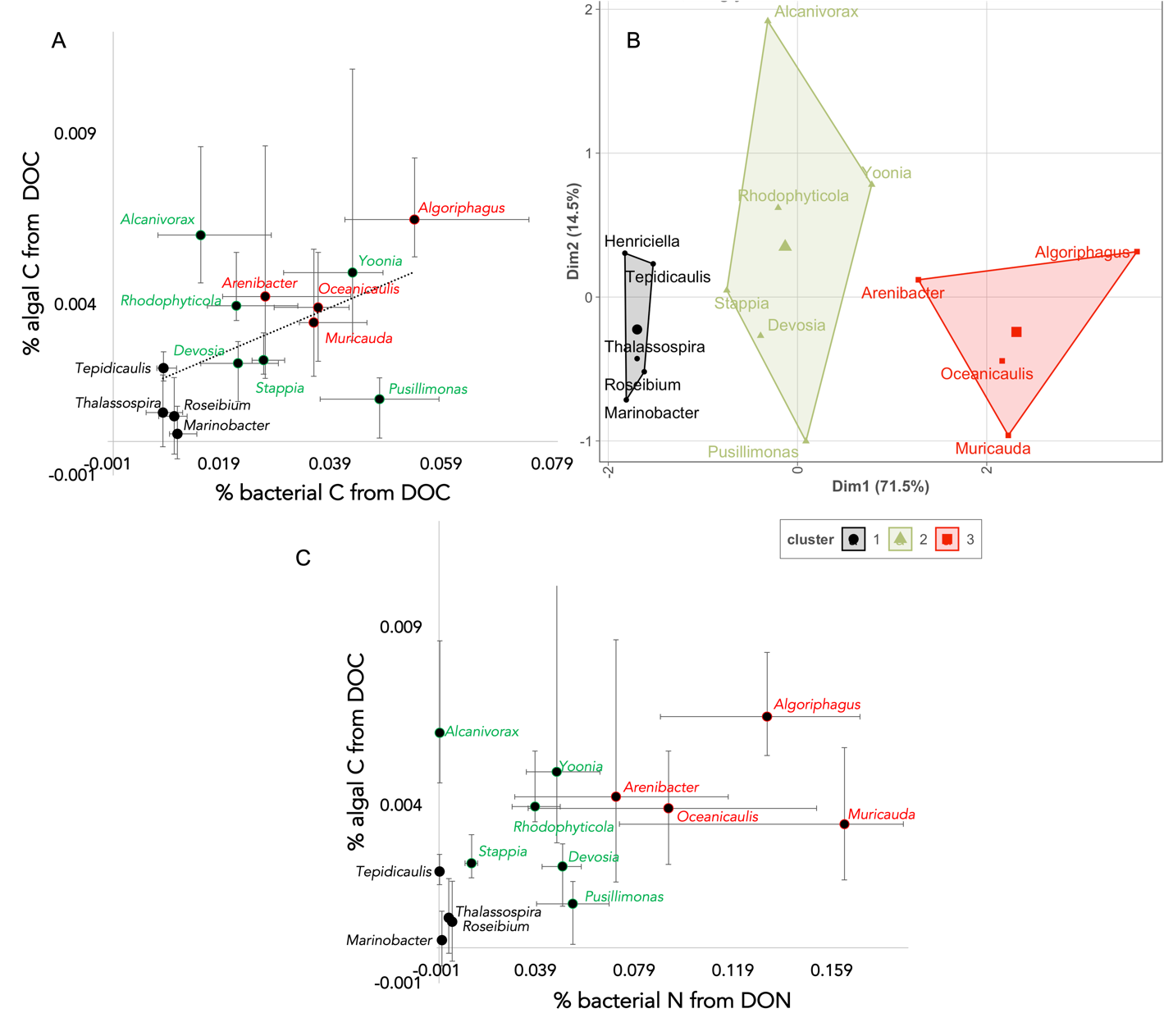

Figure S5: NanoSIMS data summarized by strain, showing median isotope incorporation of (A) bacterial carbon and algal carbon; the C and N data were used for a Kmeans clustering analysis (B), which distinguished the 15 strains into 3 categories; these 3 categories were best separated when plotting mean N incorporated by bacteria vs C remineralized to algae (C).

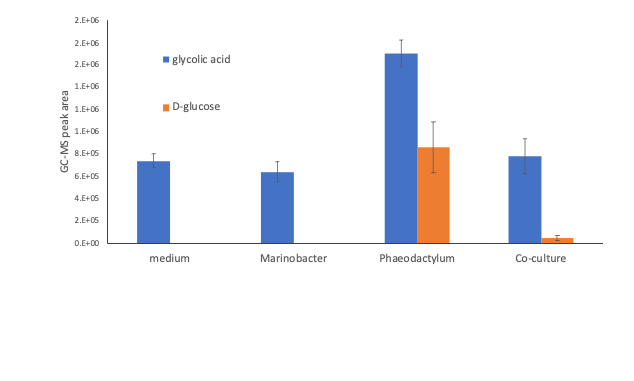

Figure S6: GC-MS detection of 2 small polar metabolites in axenic P. tricornutum, axenic Marinobacter, and co-cultures compared to media blank

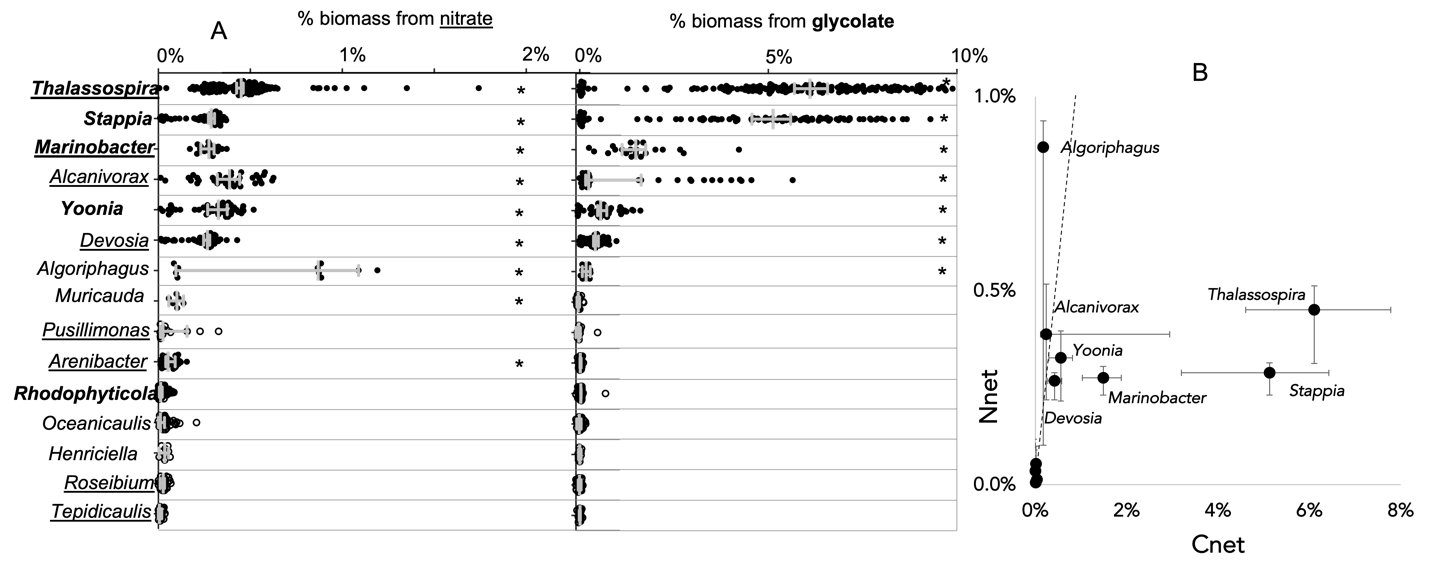

Figure S7 (A) fraction of C biomass from glycolate incorporated by single bacterial cells (median +/- 95% C.I.) as measured by 13C incorporation (solid markersand asterisks indicate cultures significantly more enriched than killed controls); (B) fraction of N biomass from nitrate incorporated by bacterial cells (median +/- 95% C.I.) as measured by 15N incorporation (solid markers and asterisks significantly enriched compared to killed controls); (C) relationship between C and N incorporation for the bacterial populations (error bars represent 25% and 75% confidence internals); genus names in bold indicate genomic capability of glycolate incorporation, and underlined indicate capability of assimilatory nitrate reduction.

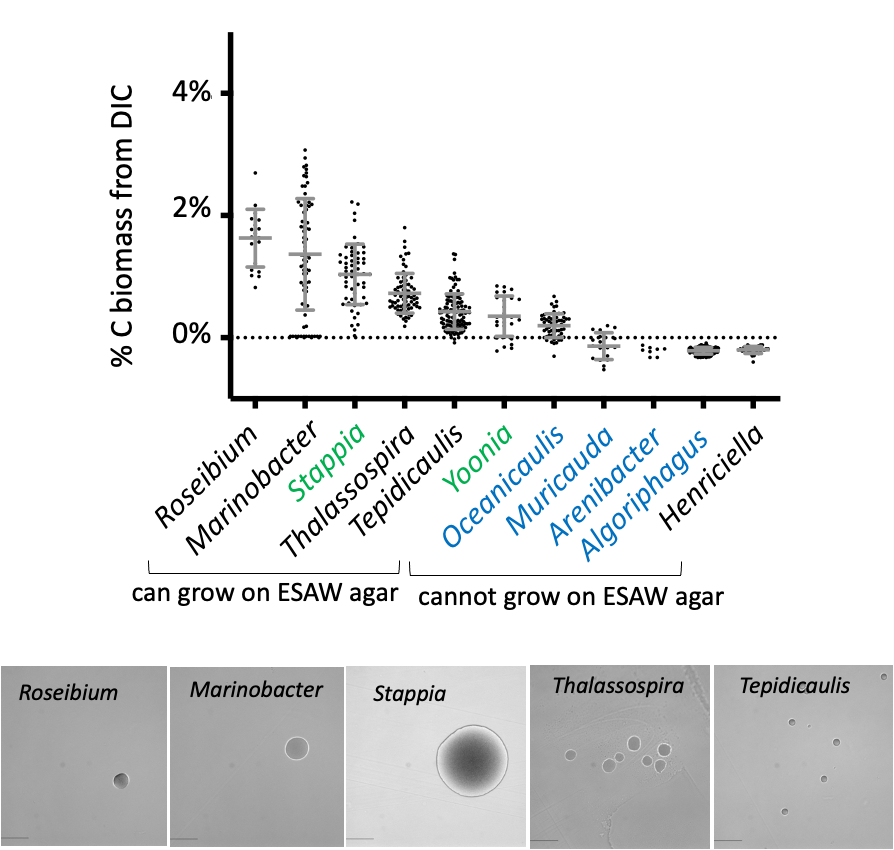

Figure S8: Net C biomass percent from inorganic carbon incorporated over a 1 week incubation by 11 bacteria grown in artificial seawater medium (ESAW). The 5 strains on the left can grow on ESAW-agar, and the 6 strains on the right cannot. Colors represent the 3 functional groups identified by Kmeans clustering of DOC and DON incorporation and remineralization.

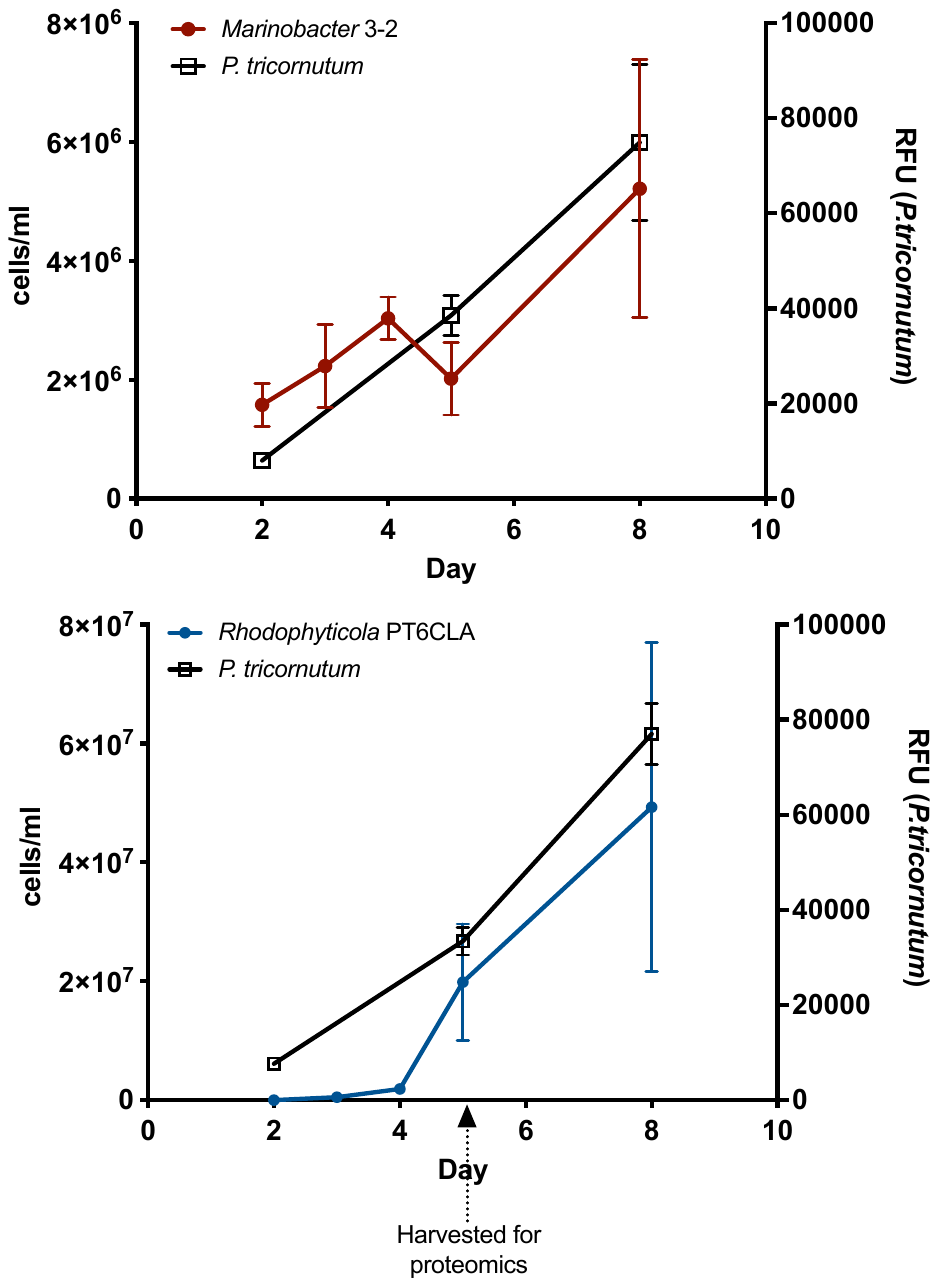

Figure S9: Averaged counts of DAPI stained bacterial cells and chlorophyll fluorescence of P. tricornutum over 8-day co-cultivation, indicating harvest for proteomics
